# In vivo CRISPR screening links NFKB1 to endocrine resistance in ER⁺ breast cancer

**DOI:** 10.1101/2025.08.11.669793

**Authors:** Cancan Lyu, Sam Hall, Mark A. Stamnes, Songhai Chen

## Abstract

Resistance to endocrine therapy (ET) remains a major clinical challenge in the treatment of estrogen receptor–positive (ER⁺) breast cancer, underscoring the need for novel therapeutic targets. To identify genetic drivers of ET resistance, we conducted an in vivo genome-wide CRISPR-Cas9 screen in MCF7 cells implanted into ovariectomized nude mice under estrogen-deprived conditions. NFKB1 emerged as a top candidate whose loss promoted estrogen-independent tumor growth and recurrence. Functional studies confirmed that NFKB1 deficiency enhanced tumorigenicity and conferred resistance to tamoxifen and fulvestrant both in vitro and in vivo. Mechanistically, transcriptomic and biochemical analyses revealed that NFKB1 loss activated canonical NF-κB signaling, leading to inflammatory gene induction and hyperactivation of ER signaling. Importantly, pharmacologic inhibition of NF-κB signaling restored ET sensitivity in NFKB1-deficient cells. Clinically, NFKB1 downregulation was enriched in ER⁺ breast tumors and associated with poor patient outcomes. Collectively, these findings establish NFKB1 as a key suppressor of ET resistance, uncover a mechanistic link between inflammation and ER reactivation, and highlight NF-κB signaling as a therapeutic vulnerability in NFKB1-deficient ER⁺ breast cancer.

## Introduction

Estrogen receptor–positive (ER+) breast cancer accounts for over 75% of all breast cancer diagnoses. Activated ER signaling is a key driver of tumor initiation and progression through its promotion of cell proliferation (1). Accordingly, endocrine therapies (ETs) that inhibit ER signaling remain the foundation of treatment for ER+ breast cancer (2). These include selective ER modulators (SERMs, e.g., tamoxifen), selective ER degraders (SERDs, e.g., fulvestrant), and aromatase inhibitors (AIs, e.g., anastrozole), with tamoxifen historically serving as a frontline agent (3, 4).

While ETs have markedly improved outcomes, therapeutic resistance remains a significant clinical challenge. Some ER+ tumors exhibit de novo resistance, failing to respond to initial ET, while over 30% of patients eventually develop acquired resistance, often leading to metastatic disease (4). Metastasis is associated with a steep decline in prognosis—reducing 5-year survival to ∼20% and median survival to just three years (5).

The mechanisms underlying ET resistance are multifactorial and have been extensively studied (4, 5). De novo resistance is often attributed to loss of ERα expression, mutations in *ESR1*, or deficiencies in drug-metabolizing enzymes like CYP2D6 (6, 7). Acquired resistance typically involves sustained proliferative signaling driven by alterations within ER signaling, cross-talk with receptor tyrosine kinases (e.g., HER2, EGFR, FGFR, IGFR), dysregulation of cell cycle regulators (e.g., cyclin D1, MYC), and activation of survival pathways such as PI3K/AKT/mTOR, MAPK, Hedgehog, and deregulated NF-κB signaling (4, 5). The aberrant NF-κB activity plays a crucial role in promoting cell survival, proliferation, and inflammation, thereby contributing to the development and maintenance of ET resistance in ER^+^ breast cancer(8). Recent therapeutic advances—such as combining ET with CDK4/6 inhibitors (e.g., palbociclib) or mTOR inhibitors (e.g., everolimus)—have provided clinical benefit but remain limited by cost, toxicity, and non-universal efficacy(5). Thus, there is a critical need to identify new therapeutic targets and predictive biomarkers to improve the management of ET-resistant ER^+^ breast cancer.

The advent of CRISPR-Cas9 genome editing has transformed functional genomics by enabling precise, high-throughput loss-of-function screening (9, 10). Genome-wide CRISPR-Cas9 screens have been instrumental in identifying key mediators of drug resistance, tumor progression, and metastasis across multiple cancer types (10). In ER^+^ breast cancer, several such in vitro screens have revealed novel regulators of endocrine therapy resistance. These include loss of C-terminal SRC kinase, which leads to increased Src signaling; loss of *NR2F2*, which impairs tumor growth and restores sensitivity to endocrine therapy; and alterations in purine biosynthesis pathways (11–13).

While these in vitro studies have provided valuable insights and identified promising biomarkers and therapeutic targets, they do not fully capture the complexity of the tumor microenvironment or the influence of systemic factors that shape treatment response in vivo (14). As a result, critical mechanisms driving endocrine resistance may remain undiscovered. Therefore, incorporating in vivo CRISPR-based screening approaches is essential to uncover additional resistance drivers and therapeutic vulnerabilities in a physiologically relevant setting (15).

In this study, we performed an in vivo genome-wide CRISPR-Cas9 screen to identify genes that modulate tamoxifen sensitivity and contribute to tumor recurrence in estrogen receptor–positive (ER⁺) breast cancer. Among the top hits, we identified NFKB1 as a critical regulator of endocrine therapy response. NFKB1 encodes a full-length p105 protein that can be proteolytically processed into p50 via KPC1-mediated ubiquitination and proteasomal degradation (16). While p50 can heterodimerize with other NF-κB subunits, such as p65, to promote NF-κB signaling and endocrine therapy resistance, both full-length p105 and p50 homodimers have been implicated in tumor-suppressive functions (17, 18). However, the contribution of NFKB1 to ER⁺ breast cancer development and treatment resistance has remained unclear.

In this study, we demonstrate that loss of NFKB1 promotes tumor growth and ER resistance by activating NF-κB signaling pathways. This activation leads to enhanced ER signaling and diminished sensitivity to tamoxifen and fulvestrant. Importantly, pharmacologic inhibition of NF-κB signaling restored tamoxifen and fulvestrant sensitivity in NFKB1-deficient cells.

These findings establish NFKB1 downregulation as a biomarker of endocrine therapy resistance and reveal a mechanistic link between inflammatory and ER pathway reactivation. Our study highlights potential therapeutic strategies to overcome ET resistance in ER⁺ breast cancer with low NFKB1 expression.

## Results

### In vivo Genome-wide CRISPR screen identified genetic determinants of ER^+^ breast cancer development

To identify genes whose loss promotes ER+ breast cancer development, we developed a novel procedure to conduct an in vivo genome-wide CRISPR knock-out screen using MCF7 cells (Figure 1A). MCF7 cells, stably expressing firefly luciferase, were transduced with the genome-scale CRISPR knock-out library (GeCKO V2) via lentiviral guide RNA (sgRNA). Transduction conditions were optimized to ensure that over 90% of surviving cells harbored a single sgRNA-expressing lentiviral integrant, maintaining greater than 95% library representation. Following infection, MCF7 cells were cultured in puromycin for two weeks to select for stable sgRNA expression and eliminate cells with essential gene knockouts for proliferation.

**Figure 1.**
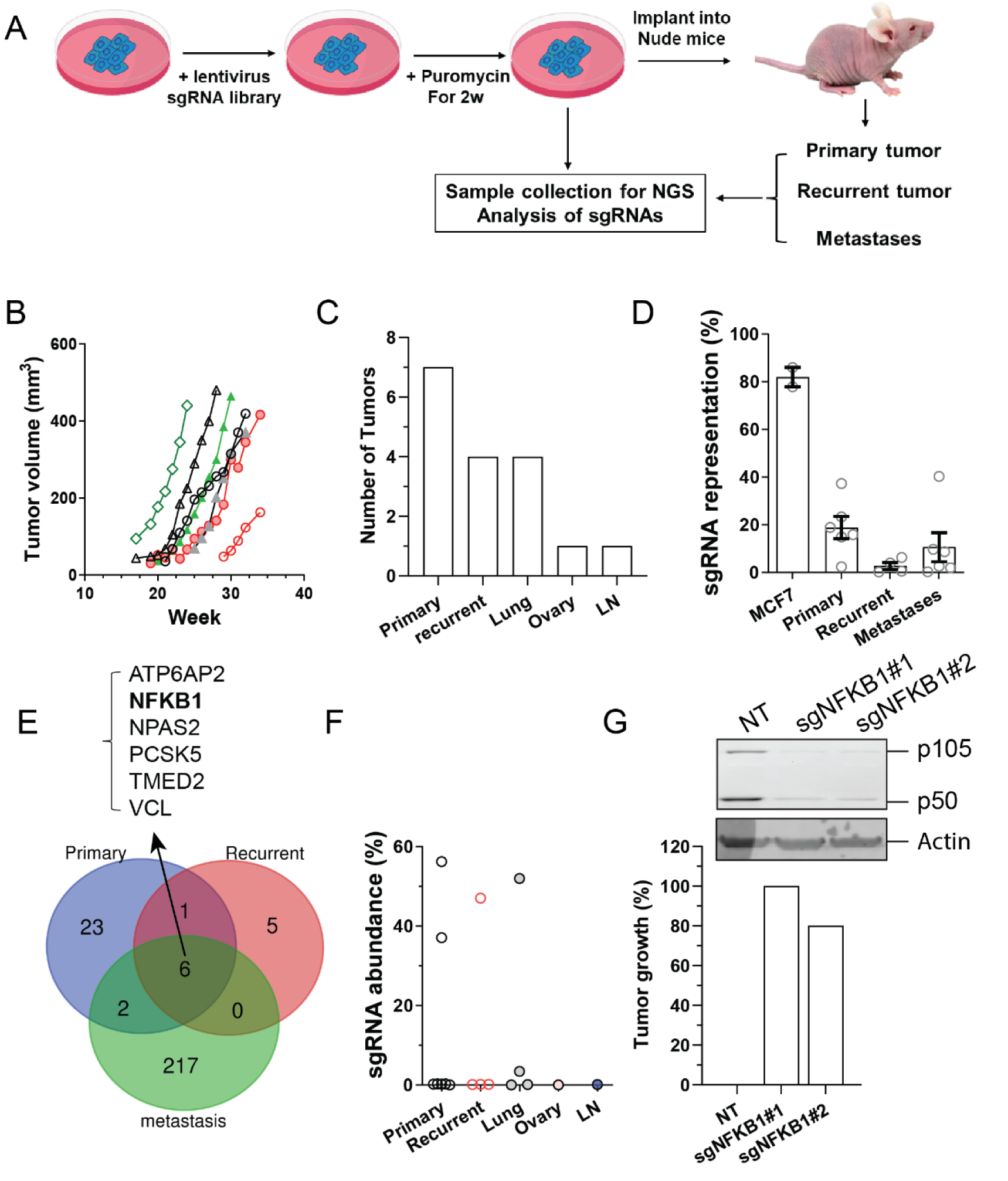
In vivo genome-wide CRISPR screen identifies NFKB1 as a key driver of estrogen-independent tumor growth in ER⁺ breast cancer. (A) Schematic of the *in vivo* genome-wide CRISPR knockout screen. MCF7-luciferase cells were transduced with the GeCKO v2 library and orthotopically implanted into ovariectomized nude mice without exogenous estrogen supplementation. **(B)** Primary tumor formation in 7 of 10 mice following implantation of GeCKO library-transduced MCF7 cells. **(C)** Overview of primary, recurrent, and metastatic tumors (including lung and ovarian metastases) identified post-resection. **(D)** sgRNA diversity and representation across different sample types: library-transduced MCF7 cells (input), primary tumors, recurrent tumors, and metastatic lesions. **(E)** Differential enrichment analysis of sgRNAs across primary, recurrent, and metastatic tumors, highlighting candidate genes, with NFKB1 identified as a top hit. **(F)** Relative enrichment of NFKB1-targeting sgRNAs (sgNFKB1) in primary, recurrent, and metastatic tumors. **(G)** Validation of NFKB1 loss promoting tumor formation *in vivo*. Mice orthotopically injected with MCF7 cells expressing control sgRNA or two independent sgNFKB1 constructs developed tumors in the absence of exogenous estrogen. **Inset:** Immunoblot showing p105 and p50 protein levels in MCF7 cells expressing the indicated sgRNAs.

A portion of these selected cells (3×107 cells) was set aside for deep sequencing to identify the integrated sgRNAs, while the remaining cells were orthotopically implanted into the mammary glands of 10 female ovariectomized nude mice, at a density of 1×107 cells per mouse (Figure 1A). To specifically identify gene inactivation conferring estrogen-independent growth, we leveraged the inherent characteristic of parental MCF7 cells, which typically require exogenous estrogen for tumor formation in nude mice(19). In this experimental setup, without exogenous estrogen, parental MCF7 cells did not form palpable primary tumors or metastatic lesions. However, 7 of the 10 mice transplanted with GeCKO library-transduced MCF7 cells developed tumors within 8 months (Fig. 1B). Following complete surgical resection, confirmed by bioluminescence imaging (BLI), 4 mice experienced recurrent tumors at the primary sites, 4 mice developed lung metastasis, and one mouse each developed ovarian and lymph node metastases (Fig. 1C).

After performing deep sequencing on sgRNAs amplified via PCR from the genomic DNA of the GeCKO-transduced MCF7 cells, primary tumors, and metastatic tumors, our analysis revealed significant changes in sgRNA representation. We observed that approximately 80% of the original sgRNAs were retained in the MCF7 cells following two weeks of puromycin selection in vitro (Figure 1D).

However, a dramatic reduction in sgRNA diversity was evident in the in vivo samples. On average, less than 20% of the original sgRNAs were present in the primary and metastatic tumors, and this figure dropped to less than 3% in the recurrent tumors (Figure 1D). This substantial decrease in sgRNA representation strongly indicates the successful enrichment and selection of highly specific sgRNAs under the in vivo conditions, particularly in the absence of exogenous estrogen.

Analyzing sgRNA abundance in the tumors revealed significant variability in the top-enriched sgRNAs across individual samples. Due to this variability and the relatively low number of tumors, standard MAGeCK analysis lacked the statistical power to identify consistent hits. Therefore, we pivoted to edgeR for a differential enrichment analysis of sgRNAs in primary, recurrent, and metastatic tumors. We defined potential hits as those consistently enriched in at least two tumor samples.

Using this criterion, we identified 32, 12, and 225 potential hits in primary, recurrent, and metastatic tumors, respectively. Notably, six hits were consistently present across all tumor types, with NFKB1 emerging as one of these key genes (Figure 1E). The sgRNA targeting NFKB1 (sgNFKB1) was highly enriched in two primary tumors, one recurrent tumor, and two lung metastases. Specifically, sgNFKB1 constituted 38% and 58% of total sgRNAs in the respective primary tumors, 44% in the recurrent tumor, and 52% and 3% in the two lung metastases (Figure 1F). This strong enrichment suggests that the loss of NFKB1 may contribute to estrogen-independent ER+ breast cancer development and metastasis.

To validate these findings, we generated MCF7 cells stably expressing either a control sgRNA or one of two distinct sgRNAs targeting NFKB1 (sgNFKB1#1 or sgNFKB1#2). We then orthotopically implanted these cells into the mammary glands of five female ovariectomized nude mice for each cell line. As expected, in the absence of exogenous estrogen, none of the mice implanted with control sgRNA-expressing MCF7 cells developed tumors. In stark contrast, all five mice implanted with sgNFKB1#1-expressing cells and four out of five mice implanted with sgNFKB1#2-expressing cells developed tumors within four months (Figure 1G). These compelling results strongly support the role of NFKB1 loss in promoting estrogen-independent tumor growth.

### NFKB1 downregulation in ER^+^ Breast Cancer and its association with poor prognosis

An analysis of The Cancer Genome Atlas (TCGA) database via cBioPortal revealed that a substantial proportion of various molecular subtypes of breast tumors exhibited decreased NFKB1 mRNA expression compared to normal tissue. The reduction ranged from 3.1% to 11.9%, notably outweighing cases with increased expression (0.3% to 5.9%) (Figure 2A).

**Figure 2.**
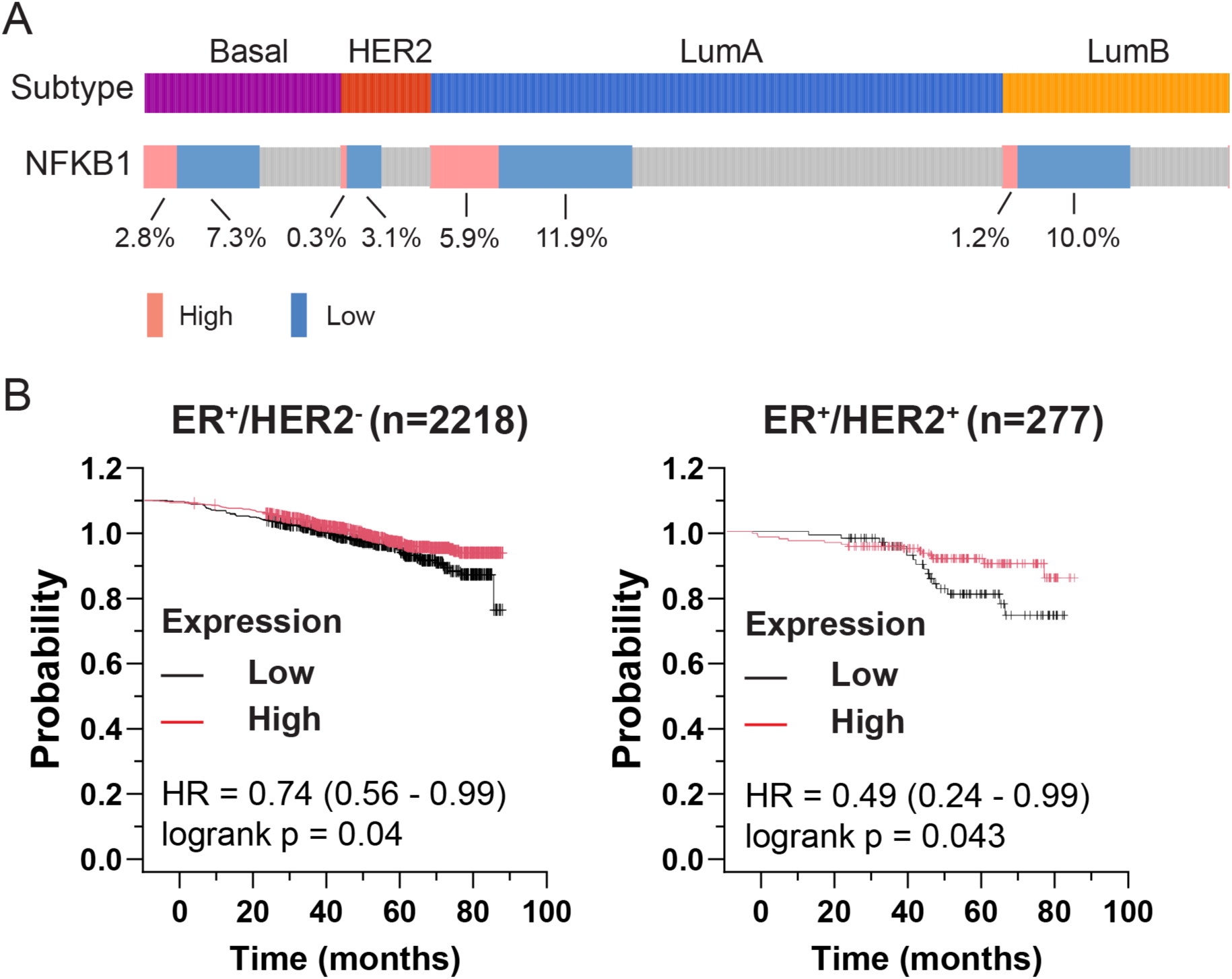
NFKB1 is frequently downregulated in ER⁺ breast tumors and correlates with poor prognosis. **(A)** Distribution of NFKB1 expression across breast cancer molecular subtypes based on TCGA data (n= 982). The frequency of tumors with high (red) and low (blue) NFKB1 mRNA expression is shown for each subtype. **(B)** Kaplan–Meier survival analysis of patients with ER⁺/HER2⁻ (left) and ER⁺/HER2⁺ (right) breast cancer stratified by NFKB1 expression levels (high vs. low). Statistical significance was determined using the log-rank test.

Specifically, ER^+^ luminal A and luminal B breast cancers displayed a higher percentage of NFKB1 reduction (11.9% and 10%, respectively) compared to other molecular subtypes such as basal (7.3%) and HER2^+^ breast cancer (3.1%) (Figure 2A). This observation suggests a particular link between NFKB1 deficiency and the development of ER^+^ breast cancer.

Furthermore, assessing the correlation between NFKB1 expression and patient survival using the Kaplan-Meier plotter demonstrated that decreased NFKB1 expression was significantly associated with poor patient outcomes in both ER^+^/HER2^-^ and ER^+^/HER2^+^ breast cancer (Figure 2B). These findings further underscore the critical importance of NFKB1 deficiency in the progression and prognosis of ER^+^ breast cancer.

### NFKB1 deficiency drives estrogen-independent growth and ET resistance in ER^+^ breast cancer

To validate the contribution of NFKB1 downregulation to estrogen-independent growth and endocrine therapy (ET) resistance in ER+ breast cancer, we generated MCF7 cells stably expressing a control shRNA or one of two distinct shRNAs targeting NFKB1 (shNFKB1#1 and shNFKB1#2). A colony formation assay performed in estrogen-depleted media demonstrated that NFKB1 downregulation significantly promoted estrogen-independent growth of MCF7 cells and conferred resistance to 4-hydroxytamoxifen (4OHT) treatment (Figure 3A-B). Consistent results were observed in T47D cells expressing shNFKB1#1 (Figure 3B). Dose-dependent analysis further revealed that NFKB1 downregulation similarly reduced the sensitivity of MCF7 cells to both 4OHT and fulvestrant. Specifically, the IC_50_ values for 4OHT and fulvestrant in control MCF7 cells were approximately 30 nM and 48 nM, respectively. In contrast, for MCF7 cells expressing shNFKB1#1 or shNFKB1#2, the IC_50_ values increased dramatically to 1-2 μM (Figure 3C-D), strongly suggesting that NFKB1 deficiency leads to broad resistance to endocrine therapies.

**Figure 3.**
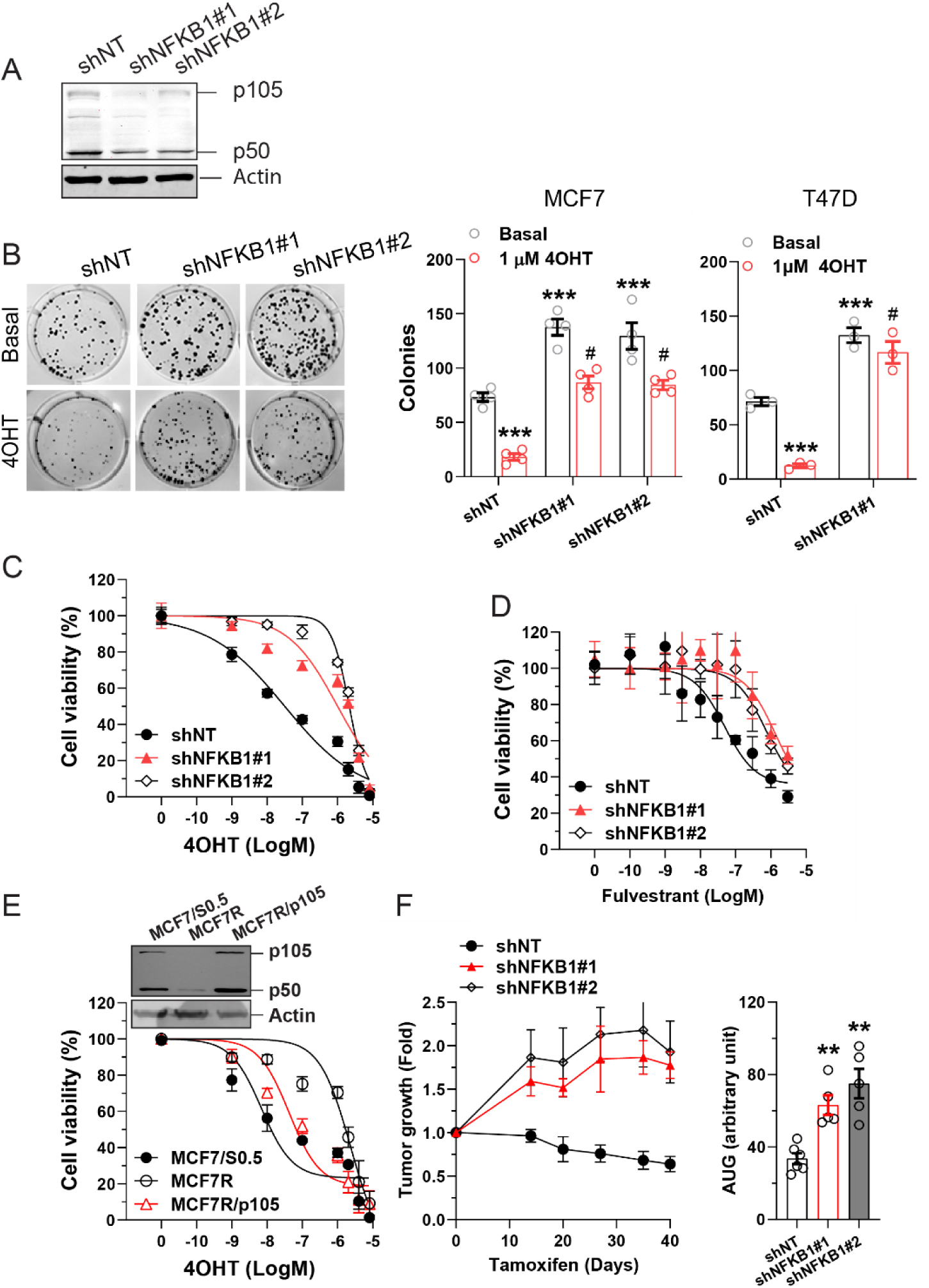
NFKB1 deficiency promotes estrogen-independent growth and endocrine therapy resistance in ER⁺ breast cancer. **(A)** Immunoblot showing p105 and p50 protein levels in MCF7 cells expressing the indicated shRNAs. **(B)** Colony formation assays under estrogen-depleted (Basal) or 1 μM 4-hydroxytamoxifen (4OHT) conditions. Left: representative images for MCF7. Right: quantification of colony formation for MCF7 and T47D cells expressing shNT or shNFKB1#1. ***p<0.001 vs shNT Basal; #p<0.001 vs shNT 4OHT; two-way ANOVA. **(C–E)** Dose–response curves showing reduced sensitivity of NFKB1-deficient MCF7 cells to 4OHT (C) and fulvestrant (D) in cell viability assays. **(E)** Dose–response curves comparing sensitivity to 4OHT in parental MCF7, 4OHT-resistant MCF7 (MCF7R), and MCF7R cells overexpressing NFKB1 (MCF7R/p105). **Inset:** immunoblot showing p105 and p50 protein levels in the indicated cell lines. **(F)** Tumor growth curves in ovariectomized nude mice implanted with MCF7 cells expressing shNT, shNFKB1#1, or shNFKB1#2. After tumor establishment with estrogen pellets, tamoxifen treatment was initiated. Tumor growth after tamoxifen treatment is shown as fold change relative to baseline. Right: quantification of tumor burden as area under the curve (AUC). **p<0.01 vs shNT; one-way ANOVA.

To further investigate the role of NFKB1 in endocrine therapy resistance, we compared its expression between experimentally established 4OHT-resistant MCF7 (MCF7R) cells and their parental counterparts. These resistant cells were generated through long-term exposure to increasing concentrations of 4OHT. Compared to parental MCF7 cells, NFKB1 expression was significantly reduced in the 4OHT-resistant MCF7 cells (Figure 3E). This reduction strongly correlated with their decreased sensitivity to 4OHT treatment, as evidenced by an IC_50_ of approximately 2.3 µM in resistant cells versus ∼6.9 nM in parental cells (Figure 3E). Notably, overexpression of NFKB1 in the resistant cells successfully restored their sensitivity to 4OHT, achieving an IC_50_ of approximately 42 nM (Figure 3F). This finding supports a critical role for NFKB1 expression in maintaining sensitivity to tamoxifen treatment.

To further validate these findings *in vivo*, we orthotopically implanted MCF7 cells expressing control shRNA, shNFKB1#1, or shNFKB1#2 into the mammary glands of ovariectomized nude mice. To facilitate tumor formation, these mice were initially implanted with silastic pellets containing 2 mg of 17β-estradiol. Once tumors reached a comparable size (approximately ∼250 mm^3^), the estrogen-containing pellets were replaced with pellets containing 1 mg of tamoxifen. As expected, the addition of tamoxifen led to tumor regression in mice implanted with control shRNA-expressing MCF7 cells (Figure 3E). In contrast, tumors derived from MCF7 cells expressing either shNFKB1#1 or shNFKB1#2 continued to grow in the presence of tamoxifen, demonstrating that NFKB1 deficiency drives both endocrine therapy resistance and sustained tumor growth *in vivo*.

### NFKB1 deficiency activates NF-κB signaling, promoting ER superactivation and ET resistance

To investigate the molecular mechanisms underlying NFKB1 deficiency-driven estrogen-independent tumor growth and endocrine therapy resistance, we performed a comprehensive transcriptomic analysis of MCF7 cells stably expressing either a control shRNA or one of two distinct shRNAs targeting NFKB1 (shNFKB1#1 or shNFKB1#2).

Differential gene expression analysis revealed that NFKB1 downregulation by shNFKB1#1 led to the upregulation of 977 genes and the downregulation of 788 genes (2.83-fold change). Similarly, shNFKB1#2 resulted in 334 upregulated and 323 downregulated genes (2-fold change), respectively (Figure 4A).

**Figure 4.**
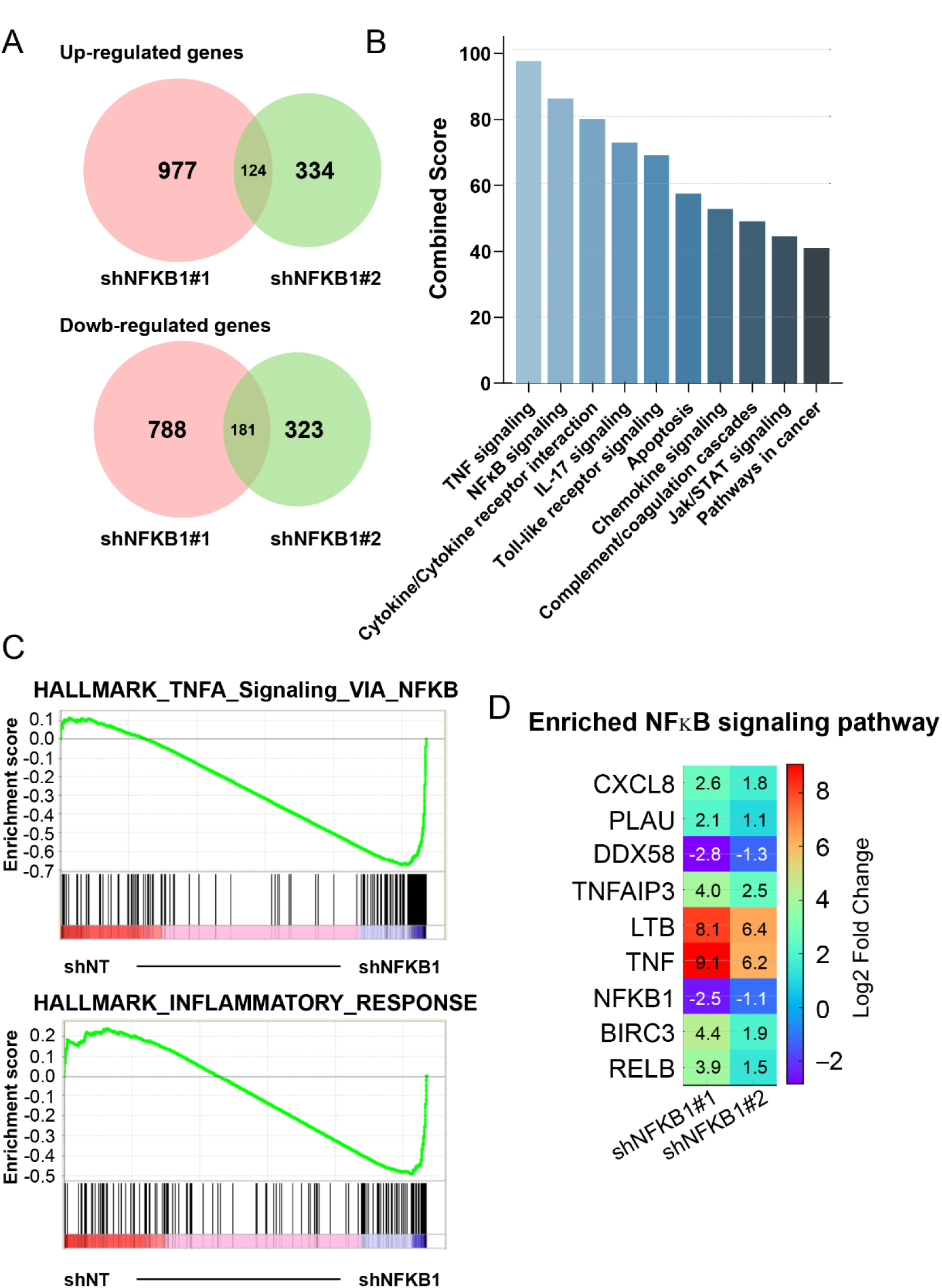
NFKB1 deficiency activates NF-κB and inflammatory signaling pathways in ER⁺ breast cancer cells. **(A)** Venn diagrams showing the number of differentially upregulated (top) and downregulated (bottom) genes in MCF7 cells expressing shNFKB1#1 and shNFKB1#2, relative shNT. Cutoffs: log₂ fold change ≥1.5 for shNFKB1#1 and ≥1.0 for shNFKB1#2; FDR < 0.05. **(B)** KEGG pathway enrichment analysis (Enrichr, combined score) of genes commonly dysregulated in both knockdown conditions. **(C)** GSEA showing significant enrichment of the TNF/NF-κB signaling pathway and inflammatory response in NFKB1-deficient cells (adjusted p-value = 0 and FDR < 0.05). **(D)** Heatmaps showing representative genes altered in the NF-κB signaling pathway (D), with log₂ fold-change values indicated for both shNFKB1#1 and shNFKB1#2. The extent of gene induction correlates with the level of NFKB1 knockdown.

Subsequent KEGG pathway enrichment analysis using Enrichr revealed that genes commonly and significantly altered by both shRNAs were strongly associated with cytokine-related inflammatory pathways. Notably, TNF and NF-κB signaling emerged as top hits enriched upon NFKB1 deficiency (Figure 4B). Gene set enrichment analysis further confirmed the upregulation of TNFα signaling via NF-κB and the inflammatory response pathway (Figure 4C). Heatmap analysis of NF-κB pathway genes showed that NFKB1 knockdown by either shNFKB1#1 or shNFKB1#2 led to significant transcriptional changes, with the extent of gene regulation corresponding to the degree of NFKB1 suppression (Figure 4D). These results suggest that NFKB1 deficiency broadly activates inflammatory signaling, which may promote estrogen-independent growth and resistance to endocrine therapy.

Quantitative PCR analysis of several downstream target genes of NF-κB revealed that NFKB1 deficiency significantly upregulated the expression of TNFα, ICAM1, PHLDA1, and CCL2, under TNF-stimulated conditions (Figure 5A). This upregulation was effectively inhibited by BMS-345541, a highly selective inhibitor of the catalytic subunits of IKK1 and IKK2(20), confirming the involvement of the NF-κB signaling pathway (Figure 5A).

**Figure 5.**
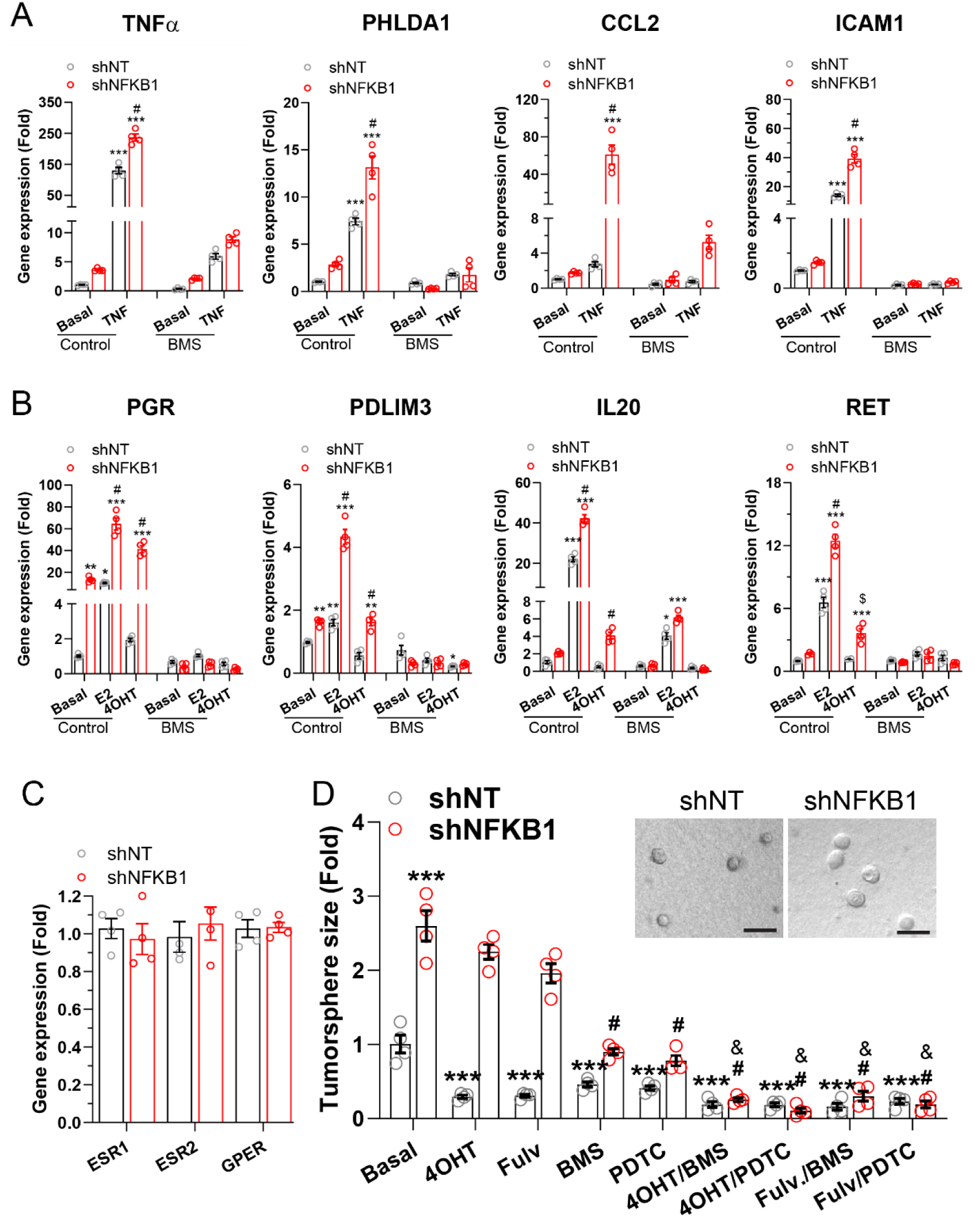
NFKB1 deficiency induces NF-κB–dependent inflammatory gene expression and estrogen receptor (ER) pathway hyperactivation. **(A-B)** Quantitative RT-PCR analysis of NF-κB target genes (A) and ER target genes (B) in MCF7 cells expressing the indicated shRNAs under basal, TNFα (100 ng/ml)-stimulated (A), or 17β-estradiol (E2; 0.1 μM)/4OHT (1 μM)-treated (B) conditions, with or without 2 μM BMS-345541 (BMS) treatment for 20 hours. *p < 0.05, **p < 0.01, ***p < 0.001 vs basal shNT; #p < 0.05 vs corresponding shNT condition; $p < 0.05 vs corresponding untreated control; two-way ANOVA. **(C)** Expression levels of ESR1, ESR2, GPER, and ESRRA in MCF7 cells expressing the indicated shRNAs. **(D)** Effects of NF-κB inhibition on estrogen-independent growth and endocrine therapy response in MCF7 cells cultured in Matrigel under estrogen-depleted conditions. Treatments: 4OHT (1 μM), fulvestrant (Fulv, 1 μM), BMS-345541 (2 μM), and pyrrolidinedithiocarbamate ammonium (PDTC, 20 μM). ***p < 0.001 vs. basal shNT; #p < 0.001 vs. basal shNFKB1; $p < 0.01 vs. corresponding 4OHT or Fulv alone; two-way ANOVA. Inset: representative images of MCF7 tumorspheres formed in Matrigel.

Subsequent analysis of estrogen receptor (ER) downstream target genes indicated that NFKB1 deficiency led to a significant upregulation of PGR, PDLIM3, IL20, and RET, without altering expression of estrogen receptors ESR1, ESR2, or GPER (Figure 5B-C). This effect was observed under basal conditions and in the presence of estrogen or 4OHT treatment. Critically, this upregulation was also abrogated by BMS-345541, implicating NF-κB signaling in NFKB1 deficiency–induced ER pathway hyperactivation and endocrine therapy resistance.

To assess whether enhanced NF-κB signaling resulting from NFKB1 deficiency contributes to endocrine therapy (ET) resistance, we examined the proliferation of MCF7 cells cultured in Matrigel to form tumorspheres, thereby better mimicking the in vivo tumor microenvironment. Under estrogen-deprived conditions, NFKB1-deficient MCF7 cells displayed markedly increased estrogen-independent growth compared with control cells (Figure 5D). This growth was largely inhibited by 4OHT or fulvestrant in control cells but was resistant to these treatments in NFKB1-deficient cells. Pharmacological inhibition of NF-κB signaling with BMS-345541 or pyrrolidinedithiocarbamate ammonium (PDTC), the latter an inhibitor of IκB phosphorylation (21), partially suppressed estrogen-independent growth in both control and NFKB1-deficient cells (Figure 5D). While NF-κB inhibitors did not significantly alter the ET sensitivity of control cells, their combination with 4OHT or fulvestrant markedly restored sensitivity in NFKB1-deficient cells, resulting in near-complete inhibition of tumorsphere formation (Figure 5D). Collectively, these findings indicate that NFKB1 deficiency promotes estrogen-independent growth and ET resistance primarily through upregulation of NF-κB signaling.

## Discussion

Endocrine therapy is the foundation of treatment for estrogen receptor–positive (ER⁺) breast cancer, yet intrinsic and acquired resistance remain major clinical obstacles (4). While several mechanisms of ET resistance have been described—including ESR1 mutations, altered drug metabolism, and activation of alternative proliferative and survival pathways—our understanding remains incomplete, particularly regarding how inflammatory signaling integrates with hormone receptor activity in the resistant state (14). In this study, we employed an in vivo genome-wide CRISPR-Cas9 screen to systematically identify genes whose loss confers estrogen-independent growth and ET resistance. This unbiased approach revealed NFKB1 as a top hit, whose deficiency drives NF-κB pathway hyperactivation, promotes ligand-independent ER signaling, and confers resistance to both tamoxifen and fulvestrant.

Our findings align with and significantly expand upon prior research implicating NF-κB signaling as a key mediator of endocrine therapy (ET) resistance (8, 18). Aberrant activation of canonical NF-κB signaling— typically via the IKK complex and subsequent nuclear translocation of RelA/p65—has been consistently shown to enhance cell survival, proliferation, epithelial-to-mesenchymal transition (EMT), and inflammation in breast cancer (8, 18). Several preclinical studies have demonstrated that NF-κB activation contribute to ET resistance. For example, NF-κB–driven IL-6 production has been shown to activate STAT3 and promote stem-like phenotypes associated with ET resistance (22, 23). A recent study further supports this by showing that NF-κB activation drives stem-like properties in ER⁺ breast tumors and correlates with ET resistance, metastasis, and poor clinical outcomes (24). Notably, inhibition of NF-κB signaling has been shown in several models to restore endocrine sensitivity and prevent tumor recurrence (8, 18, 25).

However, the role of NFKB1 itself in this context has been poorly defined. NFKB1 encodes p105, a precursor protein that undergoes proteasomal processing to generate the p50 subunit (16). This processing step is tightly regulated by KPC1 (Kip1 ubiquitination-promoting complex subunit, an E3 ubiquitin ligase that mediates the polyubiquitination of p105, enabling its partial degradation by the 26S proteasome to form p50 (16). Unlike the transcriptionally active p65:p50 heterodimers, p50 homodimers lack a transactivation domain and often act as transcriptional repressors, particularly when complexed with histone deacetylases or BCL3 (26, 27). Prior studies have shown that overexpression of KPC1 enhances p105-to-p50 conversion, resulting in accumulation of p50 homodimers and suppression of tumor growth in multiple cancer models, including glioblastoma and melanoma (28). Conversely, loss of KPC1 impairs this proteolytic processing, leading to reduced p50 production, unchecked NF-κB activation, and accelerated tumor progression (28). The full-length p105 protein can also functions as a cytoplasmic inhibitor of NF-κB by sequestering Rel subunits, thus preventing nuclear translocation and target gene activation (29). Collectively, these features position NFKB1 as a potential negative regulator of canonical NF-κB signaling.

In support of this notion, NFKB1 has been reported to possess tumor suppressor properties in several cancer types. Mouse models have shown that p50 deficiency enhances chemical carcinogenesis in the liver, and mice lacking NFKB1 developed spontaneous invasive gastric cancer and developed more alkylator-induced lymphoma than wild-type animals (30–32). In gastrointestinal tumors, loss of NFKB1 expression is associated with increased cytokine production and inflammatory response (33). Our study extends these observations to ER⁺ breast cancer, demonstrating that NFKB1 loss leads to unchecked NF-κB signaling, upregulation of inflammatory and ER target genes, and ET resistance. Notably, NFKB1 expression was significantly reduced in tamoxifen-resistant cells and its re-expression restored drug sensitivity. Analysis of breast tumor datasets revealed frequent downregulation of NFKB1 in a substantial fraction of ER⁺ primary tumors, particularly luminal A and B subtypes. Moreover, low NFKB1 levels also correlated with worse patient outcomes, supporting its role as a functional and prognostic tumor suppressor in this context. These findings align with recent reports showing significantly reduced NFKB1 expression in breast tumors compared to normal tissue, suggesting its potential as a biomarker of disease progression (34).

Mechanistically, our transcriptomic and enrichment analyses revealed robust upregulation of TNF and canonical NF-κB signaling in NFKB1-deficient cells. These findings were confirmed by gene expression analysis by PCR showing that NFKB1 loss results in enhanced expression of NF-κB targets, including TNF, ICAM1, CCL2, and PHLDA, alongside upregulation of ER-regulated genes such as PGR, PDLIM3, IL20 and RET—even in the presence of tamoxifen. These results point to a feedforward loop whereby inflammatory NF-κB signaling reactivates ER pathways, sustaining proliferative signaling in the absence of estrogen. This is consistent with prior findings that pro-inflammatory cues can enhance ligand-independent ER activity via phosphorylation or coactivator recruitment, suggesting that NFKB1 and its regulatory axis restrain oncogenic inflammation and maintain ER pathway homeostasis (35, 36).

Importantly, pharmacologic inhibition of NF-κB signaling—using IKK and IκB inhibitors—reversed ET resistance and suppressed the estrogen-independent growth of NFKB1-deficient cells. This suggests that targeting NF-κB may be a viable therapeutic strategy for overcoming resistance in tumors with low NFKB1 expression. Our findings are consistent with prior studies showing that elevated NF-κB activity is critical for tamoxifen resistance and tumor recurrence and targeting NF-κB pathway can restore tamoxifen sensitivity (18). As NF-κB inhibitors continue to advance in clinical development, particularly for inflammatory diseases and lymphomas, their repurposing for endocrine-resistant ER⁺ breast cancer represents a promising therapeutic opportunity.

While in vitro CRISPR screens have advanced our understanding of drug resistance mechanisms, they often fail to capture the full complexity of the tumor microenvironment, systemic influences, and the dynamic processes of tumor evolution and metastasis (10, 15). Our study underscores the unique value of in vivo genome-wide CRISPR screening in overcoming these limitations and uncovering physiologically relevant drivers of treatment response. Notably, NFKB1 emerged as a consistently enriched hit across primary, recurrent, and metastatic tumors, highlighting its central role in promoting estrogen-independent growth and recurrence.

In addition to NFKB1, our screen identified several other candidate tumor suppressors—ATP6AP2, NPAS2, PCSK5, TMED2, and VCL—whose loss may contribute to tumor progression. Among these, NPAS2, PCSK5, and VCL have previously been implicated in tumor suppression, although their specific roles in ER^+^ breast cancer remain unclear (37–41). These findings demonstrate the power of in vivo screening to uncover critical genes that may be overlooked in simplified in vitro systems.

In conclusion, our study identifies **N**FKB1 as a novel tumor suppressor and determinant of endocrine therapy response in ER⁺ breast cancer. Through regulation of NF-κB signaling, NFKB1 maintains repression of inflammatory and ER target gene networks, thereby sustaining ET sensitivity. Loss of NFKB1 disrupts this balance, leading to ER hyperactivation and drug resistance. These findings establish NFKB1 as both a predictive biomarker and a therapeutic vulnerability, with potential clinical implications for stratifying patients and guiding combination therapies that target inflammation in endocrine-resistant disease.

## Methods

### Sex as a biological variable

We restricted our experiments to female mice because the breast cancer model employed in this study—ER⁺ human breast cancer xenografts—is biologically and clinically relevant only to females.

### Reagents

17β-estrogen, 4OHT and charcoal-stripped FCS were from Sigma. BMS and PDTC were from MedChemExpress LLC. Antibodies for NFκB1 p105/p50 (no. 78869) was from Cell Signaling Technology; GADPH (sc-47724) from Santa Cruz Biotechnology. Phenol-red free growth factor reduced Matrigel (#356231) was from Corning.

### Cell lines

MCF7 and T47D cells were procured from ATCC (Manassas, VA, USA) and cultured in Dulbecco’s Modified Eagle Medium (DMEM) supplemented with 10 μg/ml insulin and 10% FCS. 4OHT-resistant MCF7 cells and their parental control MCF7/S0.5 cells were obtained from Sigma-Aldritch and cultured in phenol red-free DMEM/F12 supplemented with 1% FBS and 6 ng/ml insulin. Each cell line was cryopreserved at early passage numbers (less than six passages after acquisition) and utilized in experiments for up to 25 passages.

### CRISPR/Cas9 knockout lentiviral library

The human genome-wide CRISPR/Cas9 knockout pooled library GeCKO v2 A (Addgene#1000000048) was utilized for this study (42). This lentiviral library encodes Cas9 and comprises 65,383 guide RNAs (sgRNAs), targeting 19,050 human protein-coding genes with three distinct sgRNAs per gene, in addition to 1,000 non-targeting control sgRNAs.

### Lentiviral constructs

Single-guide RNAs (sgRNAs) targeting *NFKB1* (sg*NFKB1*#1: 5’-TTACCCGACCACCATGTCCT; sg*NFKB1*#2: 5’-GCTACCCGACCACCATGTCCT) and a non-targeting control sgRNA (ACGGGCGGCTATCGCTGACT) were cloned into the LentiCRISPRv2 vector for gene knockout studies. For *NFKB1* knockdown, lentiviral vectors pZIP-SFFV-RFP-V32 encoding control shRNA (shCT: 5’-aggcagaagtatgcaaagcat) or shRNAs targeting *NFKB1* (sh*NFKB1*#1 and sh*NFKB1*#2: 5’-cctttcctctactatcctgaa and 5’-ccagagtttacatctgatgat) were obtained from Transomic Technologies (Huntsville, AL, USA). Additionally, lentiviral vectors pLIX402-puromycin, enabling tetracycline-inducible expression of GFP and Flag-tagged *NFKB1*, were generated using the Gateway cloning system (Thermo Fisher Scientific, Waltham, MA, USA).

### Lentiviral production and MCF7 cell transduction

Library amplification and lentiviral production were performed following established protocols (43). The resulting lentiviruses were purified via ultracentrifugation, and their functional titers were quantified in MCF7 cells as previously described (43).

To ensure an optimal single-lentiviral integrant per cell and achieve an 800x coverage of the sgRNA library, 2×10^8^ MCF7 cells stably expressing firefly luciferase were transduced with the lentiviral library pool. Transduction was carried out at a Multiplicity of Infection (MOI) of 0.3 in the presence of 8 µg/mL polybrene, and the mixture was centrifuged at 800 x g for 2 hours at 37∘C. Two days post-transduction, cells underwent puromycin selection at 2 μg/mL for two weeks. Subsequently, 3×10^7^ transduced cells were cryopreserved at −80^∘^C for subsequent genomic DNA extraction and deep sequencing, while the remaining cells were prepared for implantation into nude mice.

Tranducction of MCF7, T47A, MCF47R cells with individual lentivirus was performed as we previously described (44).

### Mouse studies

For in vivo assessment, transduced MCF7 cells were orthotopically implanted into the mammary gland of 10 female ovariectomized nude mice (Charles River Laboratories) at a density of 1×10^7^ cells per mouse in 100 µL of PBS. Mice were monitored weekly by palpation for tumor formation in the absence of exogenous estrogen supplementation.

Upon reaching an approximate volume of 600 mm^3^, primary tumors were completely surgically excised. Successful excision was verified by bioluminescence imaging (BLI)(45). Subsequently, mice were monitored for recurrent tumor formation and metastatic spread using serial BLI. Excised tumor tissues were snap-frozen at −80∘C for subsequent genomic DNA extraction and deep sequencing.

To validate the effect of NFKB1 knockout on estrogen-independent tumor development, MCF7 cells transduced with lentivirus encoding either control sgRNA, sgNFKB1#1, or sgNFKB1#2 were orthotopically implanted into the mammary fat pad of female ovariectomized nude mice (1×10^6^ cells in 100 µL of PBS per mouse). Tumor formation was monitored by palpation.

To further assess the impact of NFKB1 deficiency, MCF7 cells expressing control shRNA, shNFKB1#1, or shNFKB1#2 were similarly implanted. To support tumor formation in these ovariectomized mice, a single silastic pellet containing 2 mg of 17β-estradiol was implanted. Once tumors reached approximately 250 mm^3^, the estradiol pellet was replaced with a silastic pellet containing 1 mg of tamoxifen to assess the effect on tumor progression. Tumor progression was subsequently monitored by caliper measurements.

### Genomic DNA extraction

Genomic DNA (gDNA) was extracted from the frozen MCF7 cell pellets and tumor powders as previously described (46). Briefly, samples were thoroughly resuspended in lysis buffer (50 mM Tris-HCl, 50 mM EDTA, 1% SDS, pH 8.0) containing 0.1 mg/mL Proteinase K and incubated overnight at 55°C. Following lysis, RNase A was added to a final concentration of 50 µg/mL and incubated for an additional 30 minutes at 37°C. To precipitate the DNA, ammonium acetate was then added to the lysate to a final concentration of 2.5 M, followed by the addition of an equal volume of cold isopropanol (or 2-2.5 volumes of cold ethanol). The mixture was centrifuged at 4,000 x g for 10 minutes to pellet the genomic DNA. The DNA pellet was subsequently washed twice with 70% ethanol. After air-drying briefly, the purified DNA was resuspended in 10 mM Tris-HCl buffer (pH 8.0) and quantified using a NanoDrop spectrophotometer (Thermo Fisher Scientific).

### sgRNA library preparation and sequencing

PCR was performed in two steps to prepare the sgRNA libraries for deep sequencing, as described previously (42). In the first step, 10 µg of genomic DNA (gDNA) were used as template to amplify lentiCRISPR sgRNAs. Each reaction had a total volume of 100 µL and utilized Herculase II Fusion DNA Polymerase (Agilent) with the following primers: F1 (AATGGACTATCATATGCTTACCGTAACTTGAAAGTATTTCG) and R1 (CTTTAGTTTGTATGTCTGTTGCTATTATGTCTACTATTCTTTCC). To ensure comprehensive library amplification from the large gDNA input, 13 separate PCR reactions were performed for each sample, and their resultant amplicons were pooled.

For the second PCR, 5 µL of the pooled first-step PCR product were used in a 100 µL reaction to attach Illumina adapters and to barcode samples. The resulting amplicons from this second PCR were gel-extracted and purified using the QIAquick Gel Extraction Kit (Qiagen). Finally, the purified libraries were sequenced on an Illumina HiSeq 3000, aiming for 20 million reads per sample.

### Bioinformatic analysis

Following Illumina sequencing, raw sgRNA counts were generated by aligning reads to the reference library via the Bowtie program (47). Due to high variability in sgRNA representation across tumor samples, standard MAGeCK analysis proved insufficiently powered to identify consistent hits. Instead, we assessed differential sgRNA enrichment using the edgeR Bioconductor package (48). To improve dispersion estimation and statistical robustness, sgRNAs with low abundance were filtered, retaining those with at least 10 counts in two or more tumor samples. Counts were then normalized using the trimmed mean of M-values (TMM) method. Differentially enriched sgRNAs were identified using a quasi-likelihood F-test, with significance defined as a false discovery rate (FDR) < 0.1 and a log₂ fold change > 1 relative to MCF7 cells.

Candidate genes were subsequently identified by collapsing enriched sgRNAs by their target gene, prioritizing those supported by multiple sgRNAs and recurrent detection in at least two tumors.

### The Cancer Genome Atlas data analyses

mRNA expression of NFKB1 in breast invasive carcinoma was analyzed using the cBioPortal for Cancer Genomics (www.cbioportal.org), specifically the TCGA PanCancer Atlas dataset. A z-score threshold of ±2 was applied to define cases with altered NFKB1 expression. Results were visualized as heatmaps stratified by molecular subtypes of breast cancer.

### Kaplan-Meier plotter analyses

The association NFKB1 mRNA expression with the survival of breast cancer patients was analyzed using the Kaplan-Meier plotter online tool (49).

### Colony formation assays

Cells were seeded at 1,000 cells per well in 6-well plates using phenol red–free media supplemented with 10% charcoal-stripped FBS as described (18). After overnight attachment, cells received indicated treatments. Media and fresh treatments were replenished every 3 to 4 days for a total of 2 weeks. Following the 2-week incubation, colonies were stained with 1% crystal violet in methanol:water (1:4) and imaged using an iBright 1500 imager (Thermo Fisher Scientific). Colony number was quantified using ImageJ software.

### Cell viability Assays

MCF7 and T47D cells were cultured for 4 days in phenol red-free media supplemented with 10% charcoal-stripped FBS. Subsequently, cells were treated with the indicated reagents for 6 days in phenol red-free media supplemented with 0.1% charcoal-stripped FBS. Cell viability was then analyzed using AlamarBlue (Thermo Fisher Scientific) according to the manufacturer’s instructions.

### Cell growth in Matrigel

Cell growth in Matrigel and treated with indicated reagents was assessed as previously described, using phenol red-free media supplemented with 0.5% charcoal-stripped FBS supplement with 10 nM hydrocortisone and 6 ng/ml insulin (50).

### Transcriptomic analysis

Total RNA was extracted from MCF7 cells cultured in complete growth media using the RNeasy micro kit (Qiagen). RNA quality was assessed via the RNA integrity number (RIN) using the RNA6000 assay (Agilent); samples with RIN > 7 were selected for sequencing. Libraries were sequenced on the BGISEQ-500 platform (BGI) as 100 bp paired-end reads with a depth of 30M reads per sample. Raw reads underwent quality control with FastQC and MultiQC, followed by adapter and low-quality read trimming using Trim Galore. Transcript quantification was performed using Salmon, and transcript-level data were aggregated to gene-level expression with Tximport, incorporating annotation. Differential gene expression was determined using DESeq2.

### Pathway Enrichment Analysis

Significantly altered genes were defined by thresholds of adjusted p-value < 0.05 and log₂ fold change ≥ 1.5 for shNFKB1#1 cells and log₂ fold change ≥ 1.0 for shNFKB1#2 cells. To identify enriched signaling pathways, gene set enrichment analysis was performed using Enrichr with the KEGG 2021 Human gene set collection. Input gene lists were separated into upregulated and downregulated groups and analyzed independently. Pathway enrichment was assessed based on combined score and adjusted p-values, utilizing the Benjamini–Hochberg correction for multiple hypothesis testing. The top enriched pathways were visualized using bar plots or heatmaps, highlighting biological processes and signaling cascades significantly associated with the observed transcriptional changes.

### Analysis of gene expression by PCR

Cells were initially cultured for 4 days in phenol red-free media supplemented with 10% charcoal-stripped FBS to deplete endogenous estrogen. Cells were then exposed to the indicated reagents for 20 hours in fresh phenol red-free media supplemented with 0.1% charcoal-stripped FBS. Total RNA was extracted from the treated cells using Trizol (Thermo Fisher Scientific). Subsequently, cDNA was synthesized from 2 μg total RNA using the High-Capacity cDNA Reverse Transcription Kit (Applied Biosystems). The expression levels of target genes were assessed via qPCR using the SensiFast SYBR No-ROX kit (Bioline USA Inc, Taunton, MA, USA). The primer sequences used were as follows: 18s (AACTTTCGATGGTAGTCGCCG, CCTTGATGTGTAGCCGTTT), CCL2 (Forward: 5’-TGTCCCAAAGAAGCTGTGATC, reverse 5’-ATTCTTGGGTTGTGGAGTGAG), PHLDA1 (Forward: 5’-CATCCACATCCACACTCTCATC, reverse 5’-CTTTCAGGCAGAGTTGGAGG), ICAM1(Forward: 5’-CAATGTGCTATTCAAACTGCCC, reverse 5’-CAGCGTAGGGTAAGGTTCTTG), TNFa (Forward: 5’-ACTTTGGAGTGATCGGCC, reverse 5’-GCTTGAGGGTTTGCTACAAC), NFKB1 (Forward: 5’-GAACCACACCCCTGCATATAG, reverse 5’-GCATTTTCCCAAGAGTCATCC), PGR (Forward: 5’-TCGCCTTAGAAAGTGCTGTC, reverse 5’-GCTTGGCTTTCATTTGGAACG), PDLIM3 (Forward: 5’-TGTGACAAATGTGGGAGTGG, reverse 5’-CTTTTGCTTGAGGTTGAGGTTG), IL20 (Forward: 5’-ACCCCTGACCATTATACTCTCC, reverse 5’-TTCTTCATTGCTTCCTCCCC), RET(Forward: 5’-GGAGAAGGCGAATTTGGAAAAG, reverse 5’-CAGGACGTTGAACTCTGACAG), ESR1 (Forward: 5’-CGACTATATGTGTCCAGCCAC, reverse 5’-CCTCTTCGGTCTTTTCGTATCC), ESR2 (Forward: 5’-GGTCGTGTGAAGGATGTAAGG, reverse 5’-TTCCCACTTCGTAACACTTCC), GPER (Forward: 5’-CATTCCAGACAGCACCGAG, reverse 5’-AGTTTAGAGACATGACGTGGC).

### Western blotting analysis

Protein lysates were prepared from cells and tumor tissues and analyzed by Western blotting using the iBright 1500 (Thermo Fisher Scientific) or Odyssey (LI-COR Biotechnology, Lincoln, NE, USA) imaging system.

### Statistics

Data were expressed as mean ± SEM. Statistical comparisons between groups were analyzed by two tail Student’s *t* test or ANOVA (*P*<0.05 was considered significant).

### Study approval

All animal studies were conducted in accordance with an IACUC-approved protocol at the University of Iowa.

## Data availability

All data generated or analyzed in this study are included in this article. The complete list of CRISPR screen results is not provided at this time, as the authors are still investigating these findings; the data will be released upon publication of the related results. The processed RNA-seq data for MCF7 cells are deposited in https://data.mendeley.com/datasets/5xym7nhskw/1. Due to export restrictions in place in China at the time of data collection, the original FASTQ files were not provided by the sequencing provider (BGI).

## Authors’ Contributions

Data curation, C. Lyu, and S. Hall; Formal analysis, C. Lyu, S. Hall, M. Stamnes and S. Chen; Methodology, C. Lyu, S. Hall; Project administration, S. Chen; Writing-original draft, S. Chen.; Writing-review & editing, S. Chen.

## Acknowledgements

This work was supported in part by NIH grant R01CA282699 and a pilot grant from Carver College of Medicine, University of Iowa.

